# Estradiol Mediates Greater Germinal Center Responses to Influenza Vaccination in Female than Male Mice

**DOI:** 10.1101/2023.11.27.568847

**Authors:** Santosh Dhakal, Han-Sol Park, Kumba Seddu, John Lee, Patrick S. Creisher, Kimberly M. Davis, Isabella R. Hernandez, Robert W. Maul, Sabra L. Klein

## Abstract

Adult females of reproductive ages develop greater antibody responses to inactivated influenza vaccine (IIV) than males. How sex, age, and sex steroid changes impact B cells and durability of IIV-induced immunity and protection over 4-months post-vaccination (mpv) was analyzed. Vaccinated adult females had greater germinal center (GC) B cell and plasmablast frequencies in lymphoid tissues, higher neutralizing antibody responses 1-4 mpv, and better protection against live H1N1 challenge than adult males. Aged mice, regardless of sex, had reduced B cell frequencies, less durable antibody responses, and inferior protection after challenge than adult mice, which correlated with diminished estradiol among aged females. To confirm that greater IIV-induced immunity was caused by sex hormones, four core genotype (FCG) mice were used, in which the testes determining gene, *Sry*, was deleted from ChrY and transferred to Chr3, to separate gonadal sex (i.e., ovaries or testes) from sex chromosome complement (i.e., XX or XY complement). Vaccinated, gonadal female FCG mice (XXF and XYF) had greater numbers of B cells, higher antiviral antibody titers, and reduced pulmonary virus titers following live H1N1 challenge than gonadal FCG males (XYM and XXM). To establish that lower estradiol concentrations cause diminished immunity, adult and aged females received either a placebo or estradiol replacement therapy prior to IIV. Estradiol replacement significantly increased IIV-induced antibody responses and reduced morbidity after the H1N1 challenge among aged females. These data highlight that estradiol is a targetable mechanism mediating greater humoral immunity following vaccination among adult females.

**Importance:** Females of reproductive ages develop greater antibody responses to influenza vaccines than males. We hypothesized that female-biased immunity and protection against influenza was mediated by estradiol signaling in B cells. Using diverse mouse models ranging from advanced age mice to transgenic mice that separate sex steroids from sex chromosome complement, those mice with greater concentrations of estradiol consistently had greater numbers of antibody producing B cells in lymphoid tissue, higher antiviral antibody titers, and greater protection against live influenza virus challenge. Treatment of aged female mice with estradiol enhanced vaccine-induced immunity and protection against disease, suggesting that estradiol signaling in B cells is critical for improved vaccine outcomes in females.

## Introduction

Human and animal studies illustrate that after receipt of either seasonal or pandemic influenza vaccines, adult females produce significantly greater quantity and quality of antibodies, which in turn provide better protection after influenza virus infection than males, at least in mice (1–6). With aging, antibody production after vaccination and protection from live influenza virus infection is reduced (3, 7, 8), with evidence that the age-associated decline in immunity is greater for females than males in response to seasonal influenza vaccines in humans (9), the pandemic monovalent 2009 H1N1 vaccine in humans (3), and universal influenza vaccine candidates in mice (10). Several studies illustrate that the effectiveness of the influenza vaccine decreases over an influenza season, likely due to waning levels of virus-specific antibodies (11–13), but whether age and sex influence the waning of influenza vaccine-induced antibody responses and protection has not been reported.

Greater vaccine-induced immunity and protection among adult females appear to be mediated by differential regulation of genes associated with B cell function. *Toll-like receptor 7 (Tlr7)* plays an important function in antibody isotype switching and antibody production in the germinal centers (GC) (14, 15). Adult female mice have a greater expression of the X-linked *Tlr7* gene in splenic B cells following vaccination as compared to adult males, with deletion of *Tlr7* eliminating sex differences in vaccine-induced immunity and protection (4). Increased DNA methylation in the promoter of *Tlr7* contributes to greater *Tlr7* expression in B cells from vaccinated female than male mice (4), but with known and putative estrogen response elements in the promoter of *Tlr7* (16), regulation of *Tlr7* expression by estrogen receptor signaling cannot be ruled out.

Expression of activation-induced cytidine deaminase (*Aicda*) mRNA, the gene that encodes activation-induced deaminase (AID) enzyme, is involved in somatic hypermutation (SHM), and shows greater expression in splenic B cells isolated from vaccinated adult females than adult male mice, with deletion of *Aicda* eliminating sex differences vaccine-induced immunity and protection (6). These data suggest that sex differences humoral immunity is dependent on greater class switch recombination and SHM in B cells from female than male mice. Regulation of these processes in B cells by sex steroids has been established in autoimmune disease mouse models (17, 18), but less so in the context of inactivated vaccines, where humoral immunity is the correlate of protection (19). Both in humans and mice, estradiol is positively, and testosterone is negatively, associated with antibody titers after influenza vaccination (2, 3). Moreover, in adult mice, sex differences in vaccine-induced immunity are eliminated by removal of the gonads and restored by exogenous sex steroid replacement in gonadectomized male and female mice (3). The contributions of gonadal sex versus sex chromosome complement to sex differences in influenza vaccine-induced immunity and protection have not been systematically investigated. Because the estrogenic changes with aging affect vaccine-induced immunity (3), we hypothesized that sex steroids more than sex chromosome complement would mediate sex differences in influenza vaccine-induced immunity and protection against infection. Whether changes in sex steroid concentrations affect numbers of antibody producing B cells, titers of antiviral antibody, or both was further explored. Finally, consideration was given to the therapeutic use of estrogen replacement therapy for improving vaccine-induced immunity and protection in aged female mice.

## Results

### Vaccinated adult females have greater numbers of antibody producing B cells, antibody responses, and protection against influenza than males, which changes with advanced age

Previous studies from our group reveal that after receipt of an inactivated 2009 H1N1 vaccine, adult females have greater neutralizing antibody responses, more cross-reactive IgG antibodies, more GC B cells in spleens, and greater SHM frequencies in regions of the recombined V genes in splenic GC B cells than males (3, 4, 6). Whether these sex differences in humoral immunity change with aging has not been explored. In draining lymph nodes (i.e., inguinal, and popliteal) (20) collected at 35 dpv (i.e., 14-days post boost) from vaccinated animals, frequencies and total numbers of GC B cells and plasmablasts were determined by flow cytometry (**Fig. 1A**). Adult mice had significantly greater frequencies and numbers of GC B cells (**Fig. 1B-C**) and plasmablasts (**Fig. 1D-E**) in their draining lymph nodes than aged mice. Adult females had significantly greater numbers of GC B cells (**Fig. 1C**) as well as greater frequencies and numbers of plasmablasts (**Fig. 1D-E**) in draining lymph nodes than adult males. Sex differences in B cell numbers and proportions were not observed in lymph nodes from aged mice (**Fig. 1B-E**). The frequencies and numbers of GC B cells were also determined at 35 dpv in the spleen. As observed in the lymph nodes, the frequencies, and numbers of GC B cells in the spleens of vaccinated mice were greater among adult than aged mice, with adult females having more GC B cells than adult males (**Fig. 1F-G**). Splenic GC B cells were sorted and the J_H_4 intronic regions of the recombined V genes were sequenced. Consistent with previous results (6), the mutation frequency in the J_H_4 intronic region showed a trend of greater frequencies in splenic GC B cells from adult females than adult males (**Fig. 1H**, *p=0.1*), with the sex difference in SHM not observed among aged animals who generally had greater variability in SHM frequencies in splenic GC B cells than among adult animals (**Fig. 1H**). These data illustrate an age-associated reduction in the numbers of GC B cells and plasmablasts, but not in SHM, with adult females having greater numbers of GC B cells and plasmablasts in lymphoid tissues than adult males, which is mitigated with aging.

**Figure 1.**
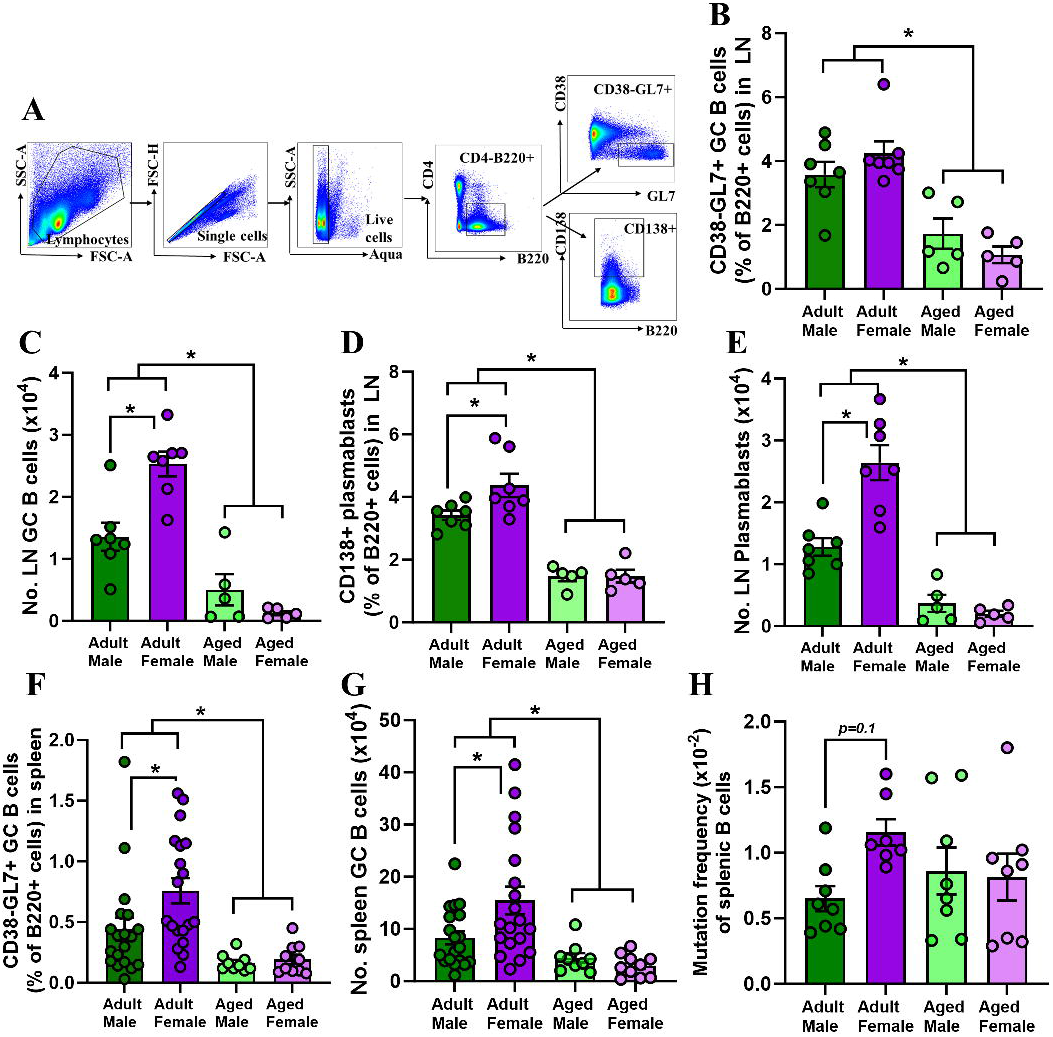
The frequency and number of germinal center (GC) B cells and plasmablasts are greater in the draining lymph nodes and spleens from vaccinated adult, but not aged, females than males. Adult (8-10 weeks old) and aged (17 months old) male and female C57BL/6CR mice were vaccinated twice with inactivated 2009 H1N1 vaccine in a 3-week interval. At 35 days post-vaccination (i.e., 14 days post-boost), draining lymph nodes and spleens were collected, single-cell suspensions were prepared, and flow cytometry was performed to measure the frequencies and numbers of GC B cells and plasmablasts. (A) Representative flow plots are shown from the lymph nodes of one adult female mouse. Frequencies and numbers of (B, C) GC B cells and (D, E) plasmablasts in the lymph nodes were quantified. The frequencies and numbers of GC B cells in the spleen were quantified (F, G), GC B cells were sorted, and (I) mutation frequency in the J_H_4 intronic region of sorted splenic GC B cells was measured. Data represent the mean ± standard error of the mean (n=5-19/group), and asterisks (*) represent significant differences (p<0.05) between the groups based on two-way ANOVAs followed by Tukey’s multiple comparisons tests in GraphPad Prism 10.1.0.

In addition to having more antibody producing cells in draining lymph nodes and spleens at 1-mpv, vaccinated adult females had greater titers of anti-2009 H1N1-specific IgG, IgG2c, and virus neutralizing, but not IgG1, antibodies than adult males **(Fig. 2A-D)**. While adult mice had greater antibody responses than aged mice, no sex differences in antibody titers were observed among aged mice (**Fig. 2A-D**). Concentrations of estradiol were greater in adult females than either males or aged females (**Fig. 2E**), reflecting the patterns observed for both antibodies producing cells and antibody titers. In contrast, adult males had greater testosterone concentrations than either females or aged males (**Fig. 2F**), which did not reflect the patterns of vaccine-induced immunity. These data suggest that numbers of antibody producing B cells and titers of antiviral antibody are greatest in the animals that have the highest circulating concentrations of estradiol.

**Figure 2.**
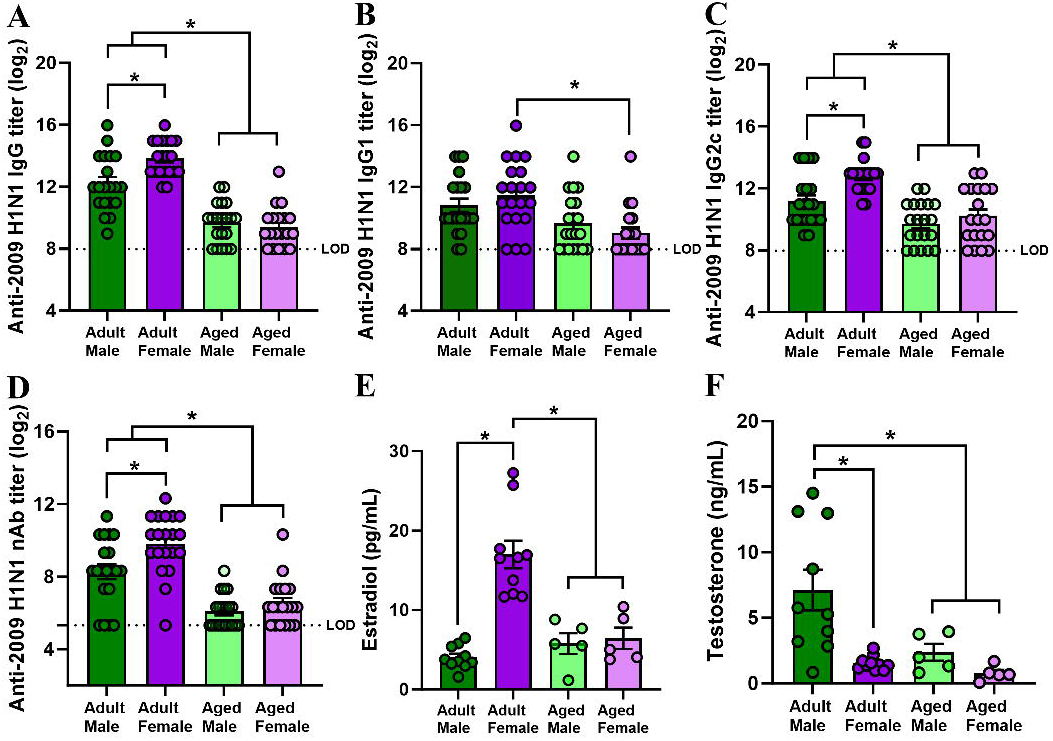
Adult, but not aged, females have higher antibody titers at 1-month post-vaccination (mpv). Adult (8-10 weeks old) and aged (17 months old) male and female C57BL/6CR mice were vaccinated twice with inactivated 2009 H1N1 vaccine in a 3-week interval. At 35 days post-vaccination (i.e., 1 mpv), plasma samples were collected to determine the titers of anti-2009 H1N1 influenza virus-specific (A) IgG, (B) IgG1, (C) IgG2c, and (D) virus-neutralizing antibody (nAb) titers and to measure the concentrations of (E) estradiol and (F) testosterone. Data represent the mean ± standard error of the mean (n=5-20/group), asterisks (*) represent significant differences (p<0.05) between the groups based on two-way ANOVAs followed by Tukey’s multiple comparisons tests in GraphPad Prism 10.1.0.

### Sex differences in vaccine-induced antiviral antibody responses are durable over time among adult, but not aged, mice

To explore sex and age differences in the durability of vaccine-induced antibody responses and protection, vaccinated adult and aged male and female mice were followed for 4-mpv, and plasma samples were collected at each month to measure anti-2009 H1N1 antibody responses. Adult mice maintained highly detectable anti-2009 H1N1 IgG, IgG2c, and nAb titers for up to 4-mpv, with females maintaining greater antibody responses than males for the duration of the study (**Fig. 3A-C**). In contrast, after 1-mpv anti-2009 H1N1 IgG, IgG2c, and nAb titers fell below the limits of assay detection among aged mice, with no sex differences observed (**Fig. 3A-C**). Vaccinated adult and aged male and female mice were challenged with a 2009 H1N1 drift variant virus at either 1 or 4 mpv. Infectious virus titers were measured in the lungs at 3 dpc and were significantly lower among adult than aged mice, with adult females having lower pulmonary virus titers than either adult males or aged males and females both at 1 and 4 mpv (**Fig. 4A, C**). Subsets of mice were followed for 14 dpc for morbidity. At 1 mpv, vaccinated aged mice lost significantly more body mass compared with adult mice, with adult males losing more body mass than adult females after challenge with 2009 H1N1 drift variant virus (**Fig. 4B**). In contrast, after live virus challenge at 4 mpv, sex differences in protection from disease were not observed either in adult or aged animals, but adult mice were still better protected from morbidity than aged mice (**Fig. 4D**). Taken together, these data suggest that the greater vaccine-induced immunity and protection against infection, but not disease, in adult female mice were durable over time, but lost with age.

**Figure 3:**
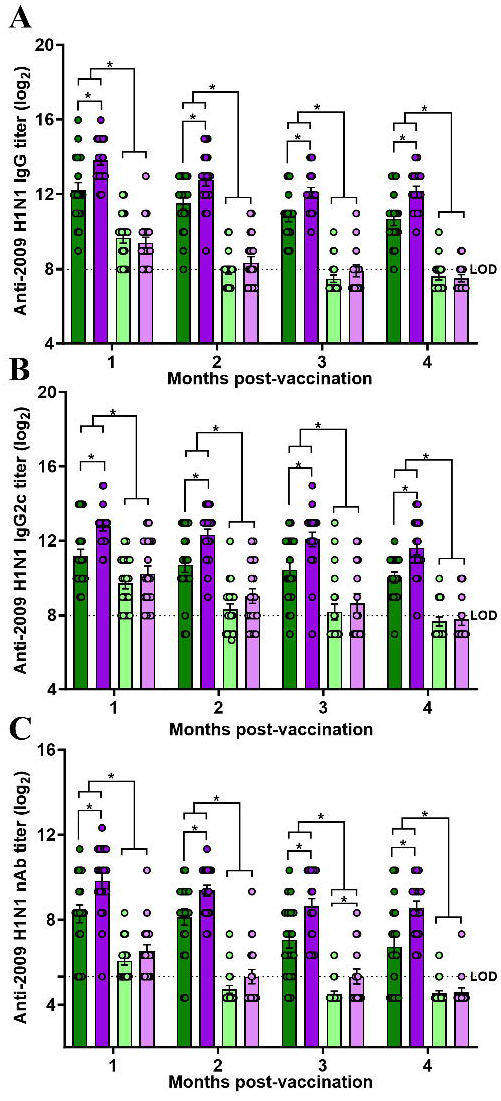
Adult female mice maintain higher titers of influenza vaccine-induced antibodies up to 4 months post-vaccination (mpv), which is mitigated with aging. Adult (8-10 weeks old) and aged (17 months old) male and female C57BL/6CR mice were vaccinated twice with inactivated 2009 H1N1 vaccine at a 3-week interval. Plasma samples were collected each month until 4 mpv and anti-2009 H1N1 influenza virus-specific (A) IgG, (B) IgG2c, and (C) virus-neutralizing antibody (nAb) titers were measured. Data represent the mean ± standard error of the mean (n=15-20/group) and significant differences between the groups are denoted by asterisks (*p<0.05) based on repeated measures two-way ANOVAs followed by Tukey’s multiple comparisons tests in GraphPad Prism 10.1.0.

**Figure 4:**
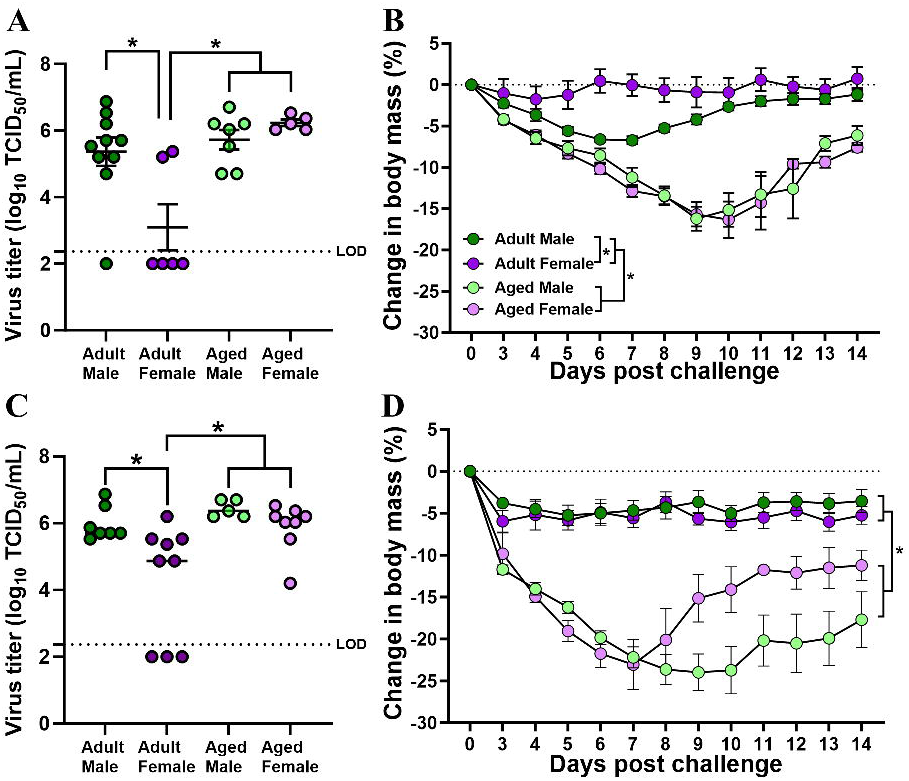
Female-biased vaccine-induced protection against infection, but not disease, is maintained for up to 4 months post-vaccination (mpv) among adult, but not aged, animals. Adult (8-10 weeks old) and aged (17 months old) male and female C57BL/6CR mice were vaccinated twice with inactivated 2009 H1N1 vaccine at a 3-week interval. At 1 or 4 mpv, vaccinated mice were challenged with 10^5^ TCID_50_ of a drift variant of the 2009H1N1 virus. (A, B) Replicating virus titers in the lungs were measured in a subset of mice at 5 days post-challenge (dpc), and (C, D) changes in body mass over a period of 14 dpc were measured in another subset of mice to compare protection from severe disease. Data represent the mean ± standard error of the mean (n=15-20/group) and significant differences between the groups are denoted by asterisks (*p<0.05) based on two-way ANOVAs or repeated measures two-way ANOVAs followed by Tukey’s multiple comparisons tests in GraphPad Prism 10.1.0.

### Sex steroids more than chromosomal complement cause sex differences in influenza vaccine-induced antibody responses and protection

Our previous work illustrated that adult females develop greater 2009 H1N1 vaccine-induced immunity and protection against 2009 H1N1 drift variant virus, which is mediated by both greater expression of the X-linked gene *Tlr7* in B cells and estrogenic enhancement of immune responses (3, 4, 6). Our current work (**Fig. 1**-**4**) also indicated that greater estradiol concentrations were associated with more durable antibody responses and protection against infection. To determine the contribution of sex steroids versus sex chromosome complement to sex differences in vaccine-induced immunity and protection, we used the FCG mouse model. The FCG mouse model involves deletion of *Sry* from ChrY and insertion of a *Sry* transgene on Chr3, resulting in: XX gonadal females (XXF), XY-gonadal females (XYF), XX*Sry* gonadal males (XXM), and XY-*Sry* gonadal males (XYM). The immunity phenotype of these FCG mice can be compared in 2x2 experimental design to separate the contribution of gonadal sex and sex steroid (i.e., testes or ovaries that produce high concentrations of androgens or estrogens, respectively) from sex chromosome complement (i.e., XX or XY) (21).

Among gonadally intact adult FCG mice, estradiol concentrations were greater in gonadal females (XXF and XYF) than gonadal males (**Fig. 5A**) and testosterone concentrations were greater in gonadal males (XYM and XXM) than gonadal females (**Fig. 5B**). At 28 dpv, we measured IgG and IgG2c binding to 2009 H1N1 as well as neutralizing antibody responses against the vaccine virus and observed that gonadal females (XXF and XYF) produce significantly greater antibody titers than gonadal males (XYM and XXM) (**Fig. 5C-E**). At 35 dpv, gonadal females (XXF and XYF) also had greater numbers of GC B cells and plasmablasts in the draining lymph nodes than gonadal males (XYM and XXM) (**Fig. 5F-G**).

**Figure 5:**
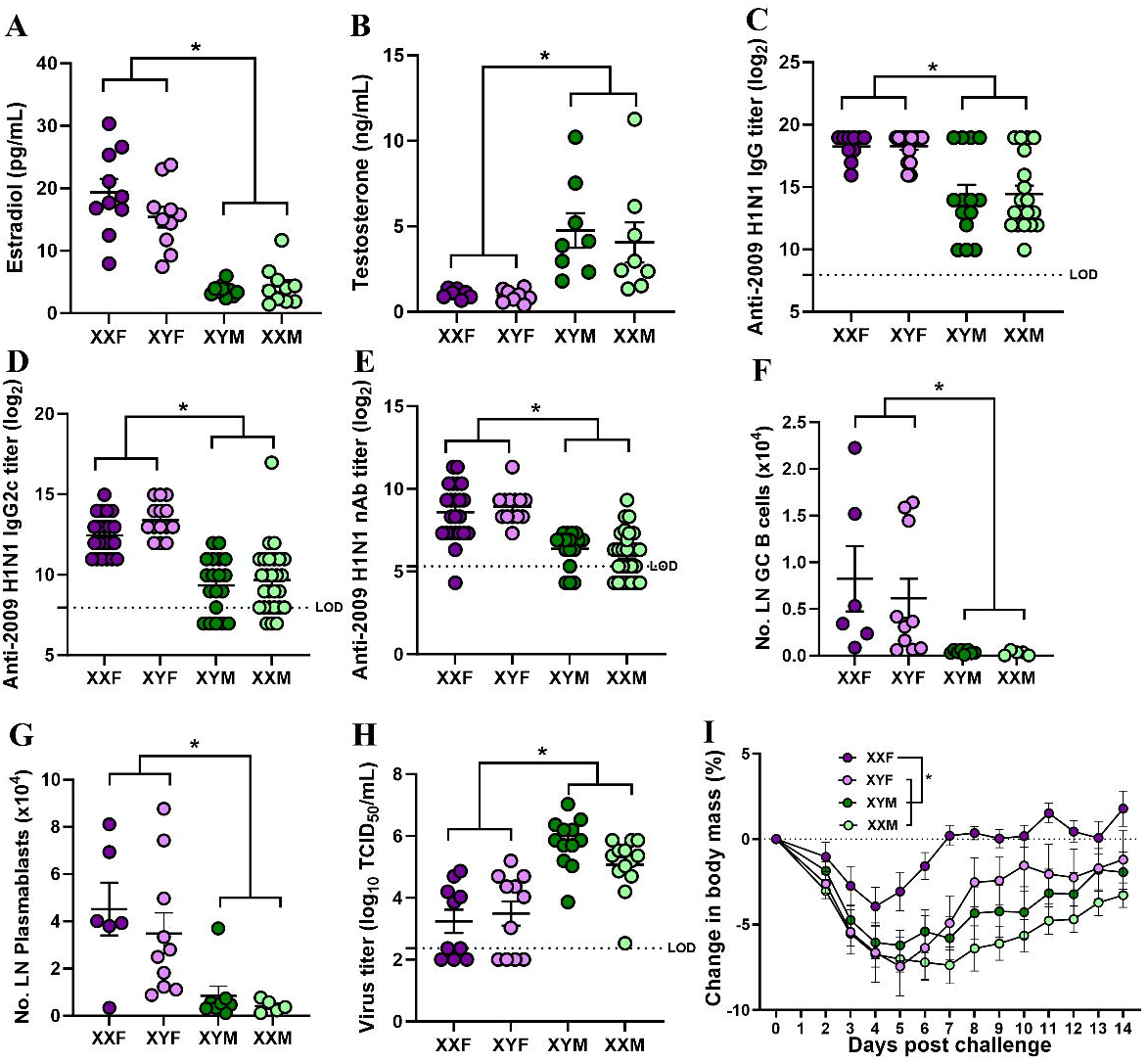
Gonadal sex more than sex chromosomal complement mediates influenza vaccine-induced immunity and protection. Eight to ten-week-old four core genotype (FCG) C57BL/6J mice were vaccinated twice with inactivated 2009 H1N1 vaccine at a 3-week interval. Plasma samples were collected at 28 days post-vaccination (dpv) and concentrations of (A) estradiol and (B) testosterone along with 2009 H1N1 influenza virus-specific (C) IgG, (D) IgG2c, and (E) virus-neutralizing antibody (nAb) titers were measured. At 35 dpv (i.e., 14 days post-boost), popliteal and inguinal lymph nodes were collected, single-cell suspensions were prepared, and the numbers of (F) germinal center (GC) B cells and (G) plasmablasts were quantified using flow cytometry. At 42 days post-vaccination (dpv), mice were challenged with 10^5^ TCID_50_ of a drift variant of the 2009 H1N1 virus and (H) replicating virus titers in the lungs were measured in a subset of mice at 5 days post-challenge (dpc) and (I) the percentage change in body mass over a period of 14 dpc was measured in another subset of mice to evaluate protection from severe disease. Data represent the mean ± standard error of the mean (n=5-27/group) and significant differences between the groups are denoted by asterisks (*p<0.05) based on two-way ANOVAs or repeated measures two-way ANOVAs followed by Tukey’s multiple comparisons tests in GraphPad Prism 10.1.0.

Vaccinated FCG mice were challenged with a drift variant of 2009 H1N1 virus, and five days later euthanized to extract lungs to measure pulmonary titers of virus. Gonadal females (XXF and XYF) had lower pulmonary titers of virus than gonadal males (XYM and XXM) (**Fig. 5H**). A separate cohort of vaccinated and infected FCG mice was followed for morbidity (i.e., mass loss after infection) as a measure of how well vaccination protected not only against infection but also disease. Vaccine-induced protection against disease revealed a gonadal sex by sex chromosome complement interaction, in which while gonadal males experienced greater disease than gonadal females, among gonadal females, XXF mice suffered significantly less morbidity than XYF mice (**Fig. 5I**). Taken together, these data suggest that sex steroids have a greater effect on vaccine-induced antibody producing B cells and protection against infection than sex chromosome complement.

### Estradiol supplementation in aged females improves influenza vaccine-induced antibody response and protection

Both the aging and FCG models illustrated that sex steroids are critical regulators of vaccine-induced humoral immunity and protection against infection with influenza virus. If reduced estradiol concentrations, in particular, cause worse vaccine-induced immunity and protection, then estradiol supplementation in aged females might rescue immunity by improving antibody responses after vaccination and protection against infection. To test this hypothesis, adult and aged female mice were implanted either with placebo or estradiol-filled capsules and vaccinated with inactivated 2009 H1N1 vaccine. At 35 dpv, IgG and IgG2c binding to 2009 H1N1 as well as neutralizing antibody responses against the vaccine virus were measured. Estradiol supplementation in aged females significantly improved anti-2009 H1N1 IgG, IgG2c, and neutralizing antibody responses after vaccination (**Fig. 6A-C**). Specifically, vaccinated aged females with estradiol produced antiviral antibody responses that were comparable to adult females with either endogenous (i.e., placebo) or exogenous estradiol and were greater than aged females that received placebo treatment. Vaccinated adult and aged female mice were challenged with 2009 H1N1 drift variant virus at 42 dpv, and infectious virus titers were measured in the lungs at 3 dpc. Estradiol supplementation in aged female mice did not significantly reduce replicating virus titers in the lungs of vaccinated mice as compared with aged females that received placebo treatment (**Fig. 6D**). In contrast, among mice that were followed for 14 dpc for morbidity, estradiol supplementation significantly reduced infection-induced morbidity as compared with placebo treatment in aged female mice (**Fig. 6E**). Vaccinated aged females treated with estradiol were as protected against severe influenza disease as vaccinated adult females that had either endogenous or exogenous estradiol. Taken together, these data highlight that estradiol replacement improves vaccine-induced antibody responses and reduces the burden of disease, but not virus replication, after infection in aged female mice.

**Figure 6:**
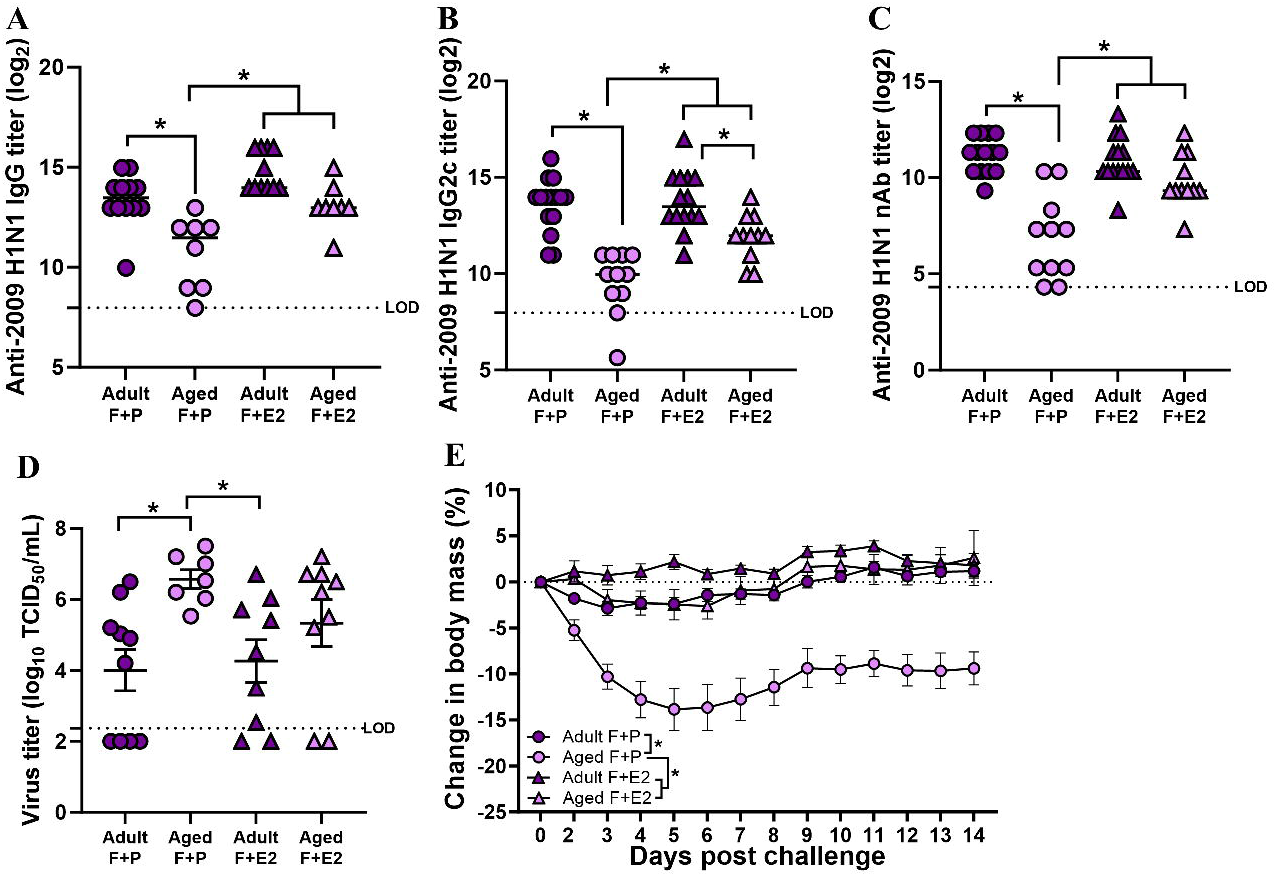
Estradiol replacement improves influenza vaccine-induced antibody responses and protection in aged female mice. Adult (8-10 weeks old) or aged (17 months old) female C57BL/6CR mice were subcutaneously implanted either with a placebo or estradiol (E2)-loaded capsules. One week after capsule implantation, mice were vaccinated with inactivated 2009 H1N1 vaccine and boosted after 3 weeks. At 35 days post-vaccination (dpv), plasma samples were collected and anti-2009 H1N1 influenza virus-specific (A) IgG, (B) IgG2c, and (C) virus-neutralizing antibody (nAb) titers were measured. At 42 dpv, vaccinated mice were challenged with 10^5^ TCID_50_ of a drift variant of the 2009 H1N1 virus. (D) Replicating virus titers in the lungs were measured in a subset of mice at 3 days post-challenge (dpc) and (E) changes in body mass over a period of 14 dpc were measured in another subset of mice to compare protection from severe disease. Data represent the mean ± standard error of the mean (n=7-15/group) and significant differences between the groups are denoted by asterisks (*p<0.05) based on two-way ANOVAs or repeated measures two-way ANOVAs followed by Tukey’s multiple comparisons tests in GraphPad Prism 10.1.0.

## Discussion

Using diverse mouse models and hormone replacement, we explored how sex and aging impact the cellular mechanisms and durability of immunity and protection to inactivated influenza vaccine (IIV). Following vaccination, greater numbers and frequencies of GC B cells and plasmablasts, as well as antiviral antibody responses are associated with better protection against infection and disease following influenza virus challenge. The novelty of our work is that we show that elevated estradiol concentrations more than other biological factors are a strong predictor of better B cell-mediated immunity and protection against infection in females as compared with males. Loss of estradiol either through aging or through use of transgenic mice significantly impairs B cell immunity and long-term protection against influenza infection and disease.

Both aged males and females have lower numbers of plasmablasts and GC B cells, less durable antibody responses, and reduced protection against both infection and disease following live virus challenge as compared with adult mice. Due to the waning of antibodies, influenza virus vaccine effectiveness declines significantly even within the same influenza season (11, 22). Such antibody waning after influenza virus infection or vaccination is more prominent among older than younger adults (13, 23). The age-associated decline in antibody responses can be broadly attributed to geriatric immunosenescence, with several B cell-specific defects associated (7, 24). After vaccination, activated B cells undergo rapid proliferation and differentiation in the GCs within the secondary lymphoid tissues, including the spleen and lymph nodes (25). SHMs and class switch recombination (CSR) occurs within the GC and together underly the production of high-affinity class-switched antibodies (26). Reduced serum antibody titers are observed among older compared to younger individuals after receipt of seasonal influenza vaccination, which is associated with lower numbers of plasmablasts (27). In humans receiving seasonal influenza vaccination, aged individuals have reduced SHM of plasmablasts as compared with younger aged individuals that results in an inability to mount antibody responses to the drifted epitopes of influenza virus (28). Reduced SHM was not observed with aging in our mice, which might reflect species-specific differences or kinetic differences in the timing of sample collection.

Sex chromosome complement (i.e., having XX or XY) can directly cause sex differences in a phenotype (e.g., humoral immunity) through an imbalance in the expression of X and Y genes that can affect immunity (29). For example, *Tlr7* is encoded on the X chromosome, can escape X inactivation in immune cells from females (30), and has greater expression in B cells from females than males following influenza vaccination (4). Sex chromosome complement also can indirectly cause sex differences in a phenotype by altering concentrations of sex steroids that can bind to nuclear receptors in immune cells to transcriptionally regulate immune cell function (31). For example, elevated testosterone in males dampens inflammatory (32) and antibody responses (2) to alter the outcome of influenza virus infection and vaccination. There can also be combined effects of genes and hormones; some X-linked genes, e.g., *Tlr7*, contain estrogen response elements and their expression can be regulated by sex steroids (16). Using the FCG mouse model, we explored whether sex differences in humoral immunity to IIV is caused by direct effects of sex chromosome complement, effects of sex steroids on immune cell function, or both. While gonadal sex, especially production of higher levels of estradiol, in adult females mediated higher levels of vaccine-induced antibody production and pulmonary virus clearance, combination of gonadal sex and sex chromosome complement appeared to modulate protection from severe disease, at least in females.

We showed that estradiol treatment can improve IIV-induced antibody production in aged females. Previous studies illustrate that estradiol treatment increases antibody response and protects female mice from influenza virus infection and these effects are mediated through ERα signaling (33). In a postmenopausal mouse model, estradiol treatment restored antibody production after vaccination with an inactivated influenza virus split vaccine (34). Because B cells have estrogen receptors, estrogens, including 17β-estradiol, can transcriptionally regulate cellular activity and function (35), in part by binding to estrogen response elements in the promoter region of estrogen-responsive genes, such as *Aicda* and directly activating AID transcription resulting in increased CSR and SHM (36, 37). In contrast, testosterone suppresses splenic B cells function by dowregulating BAFF, which is a cytokine essential for survival of splenic B cells (38). Greater serum testosterone concentrations also are associated with reduced antibody response during malaria vaccination (39).

Estradiol treatment in aged females was able to improve disease outcomes, but not virus replication, after influenza virus infection, indicating that estradiol treatment can rescue some, but not all, aspects of age-associated reductions in IIV-induced immunity. The inability of estradiol treatment to improve pulmonary virus clearance in aged mice is likely associated with age-specific changes in the pulmonary integrity and function, which are irreversible with hormone treatment. For example, in aged female mice, influenza virus infection-induced inflammation promotes fibrosis in a greater extent than in adult female mice (40). Aged female mice also have neutrophils in the lungs with altered chemotactic gene expression and tissue localization, and lymphocytes with impaired effector and memory functions as compared with adult females (40).

Overall, our study highlights that estradiol is a biological factor contributing to improved outcomes to IIV vaccine. Future studies must consider how to harness this for adjuvants or other treatments to improve vaccine outcomes in post-menopausal women. Future studies also must consider the mechanisms by which estrogens and even androgens alter the activity of B cells to impact antibody responses, which are the primary correlate of protection from influenza. Consistent observations of sex-specific effects of aging on antibody responses, we are now showing sex-specific effects of aging on numbers of antibody producing cells, including GC B cells and plasmablasts, which should be considered during the design and dosing of seasonal and universal influenza vaccines.

Future studies should explore the role of sex steroids in genetic and epigenetic regulation of GC B cell and plasmablast activity. Although AID enzyme activity was not measured in this study, the observation that IIV-specific IgG2c, but not IgG1, was greater among adult females than males and regulated by gonadal steroids highlights a fundamental role of biological sex differences in CSR. Future studies will need to consider the differential effects of gonadal steroids on the kinetics of the secretory functions of GC B cells and plasmablasts and B cell proliferation.

In humans, prior immunity caused by previous exposure to influenza viruses through infection or vaccination plays an important role in determining immune responses after subsequent influenza vaccination (41, 42). In the current study, influenza-naïve mice were used, which does not incorporate the impact of pre-existing immunity on sex or age differences in vaccine effectiveness. High-dose or adjuvanted vaccines are recommended in aged individuals for influenza and we and others have shown that females maintain greater season to season antibody durability than males among individuals 75+ years of age (9). Whether high-dose or adjuvanted vaccines could overcome the deficiency in GC B cells and plasmablasts numbers and functions in individuals with lower circulating estrogens should be explored.

## Materials and Methods

### Mice

Adult (8-10 weeks old) male and female C57BL/6CR mice were purchased from Charles River Laboratories (Frederick, MD) while the aged (17 months old) mice, originating from the Charles River Laboratories, were obtained from the National Institute on Aging (NIA). Dr. Arthur P. Arnold gifted breeder males for the FCG mouse model from the University of California, Los Angeles (43). The FCG mouse colony was maintained in-house by mating XY^-^ males with wild-type C57BL/6J females purchased from the Jackson Laboratory (Bar Harbor, ME). Genotypes were determined at weaning (i.e., at 3 weeks) by PCR analysis for the presence or absence of the *Sry* gene as described (44). Pups of the same genotype were housed together and were used at 8-10 weeks of age. Mice were housed 5/cage under standard biosafety level (BSL)-2 conditions in the Johns Hopkins Bloomberg School of Public Health animal facility with ad libitum food and water. All animal procedures were approved by the Johns Hopkins University Animal Care and Use Committee (MO20H236).

### Vaccination, challenge, and morbidity measurement

Mice were vaccinated twice, at 3-week intervals, with 20µg of mouse-adapted A/California/04/09 H1N1 (ma2009 H1N1) inactivated vaccine through the intramuscular route in the right thigh muscle (3, 4, 6). Blood samples were collected at different time points after vaccination through the retroorbital route under isoflurane anesthesia. Vaccinated mice were challenged with 10^5^ TCID_50_ of a mouse-adapted A/California/04/09 H1N1 drift variant virus (ma2009 H1N1dv) through the intranasal route under ketamine-xylazine anesthesia (3, 4, 6). To measure morbidity, body mass of the virus-challenged animals was recorded daily for a period of 14 days post-challenge (dpc).

### Hormone supplement

For estradiol supplement, adult (8-10 weeks old) or aged (17 months old) female C57BL/6CR mice were implanted subcutaneously either with an empty silastic capsule (i.e., placebo) or with a capsule loaded with17β-estradiol (5mm long), prepared as described (3).

### Antibody measurements

The levels of anti-2009 H1N1 IgG, IgG1, and IgG2c antibodies in plasma samples collected at different time points after vaccination were measured using our in-house enzyme-linked immunosorbent assays (ELISAs) (3, 4, 6). Briefly, plates were coated with 50µL/well of sodium carbonate and sodium bicarbonate coating buffer containing 2µg/mL of mouse-adapted 2009 H1N1 whole virus protein and were incubated overnight at 4°C. Next day, plates were washed 3-times, blocked with 10% skim milk solution for 1h at 37°C, and then serially diluted plasma samples were added. After 1hr incubation at 37°C, plates were washed and horse-radish peroxidase (HRP)-conjugated secondary IgG (Invitrogen), IgG1 (Invitrogen), and IgG2c antibodies (Invitrogen) were added. After 1hr incubation at 37°C, plates were washed and reactions were developed using 3,3’,5,5’-tetramethylbenzidine (TMB, BD Biosciences) for 20min, stopped using 1 N hydrochloric acid (HCL). Plates were read at 450nm wavelength using the ELISA plate reader (Molecular Devices) and the endpoint titer was calculated as the highest serum dilution with an average optical density (OD) value greater than 3-times the average OD of negative controls. Likewise, the virus-neutralizing antibody (nAb) titers on plasma samples, against the vaccine virus (i.e., ma2009 H1N1 virus), were measured using a Madin-Darby canine kidney (MDCK) cells-based microneutralization assay, as previously described (3, 4, 6).

### Sex steroid measurement

Concentrations of sex steroids on plasma samples were measured using commercial testosterone (IBL America, Minneapolis, MN) and estradiol (Calbiotech Inc., El Cajon, CA) ELISA kits, as per the manufacturer’s instructions (3, 10).

### Virus titration in lungs

For virus titration, lung samples collected at 3 or 5-dpc were homogenized, lung-homogenates were 10-fold serially diluted in serum-free media and then transferred in six replicates in 96-well cell culture plates confluent with MDCK cells. Plates were incubated for 6 days at 32°C followed by fixation with 4% formaldehyde, staining with naphthol blue-black solution, and virus titer calculation by Reed and Muench method (6, 10).

### Flow cytometry

The number of GC B cells (CD4^-^B220^+^CD38^-^GL7^+^) and plasmablasts (CD4^-^B220^+^CD138^+^) in the lymph nodes (i.e., mix of popliteal and inguinal) or spleens collected at 35dpv were determined using flow cytometry (6). Antibodies used were PerCP-cy5.5 rat anti-mouse CD4 (#55095, clone: RM4-5, BD Biosciences), PE-Cy7 rat anti-mouse CD45R/B220 (#552772, clone RA3-6B2, BD Biosciences), BV421 rat anti-mouse CD38 (#562768, clone 90/CD38, BD Biosciences), FITC rat anti-mouse T-and B-cell activation antigen (clone GL7, #553666, BD Biosciences), and APC rat anti-mouse CD138 (#558626, clone 281-2, BD Biosciences). Cells were acquired using the LSR II instrument (BD Biosciences) and analyzed using FlowJo software v.10.8.1 (BD Life Sciences).

### Somatic hypermutation (SHM)

For SHM, splenic GC B cells (B220^+^CD38^-^ GL7^+^) were sorted at 35dpv using BD FACS Aria Fusion (BD Biosciences). Sorted cells were then lysed in a digestion buffer, genomic DNA was isolated by phenol/chloroform extraction and ethanol precipitation, and the J_H_4 intronic region was amplified using a nested polymerase chain reaction (PCR) protocol. The J_H_4 intronic DNA (492 bp) was sequenced and mutations in the unique VDJ clones were analyzed as described earlier (6).

### Statistical analysis

Data were analyzed in GraphPad Prism version 10.1.0. Sex steroids concentration, antibody titers, virus titers in the lungs, numbers of GC B cells and plasmablasts, and SHM frequencies were compared using two-way ANOVA followed by Tukey’s multiple comparisons. Antibody responses up to 4-mpv and change in body mass after virus challenge were compared using repeated measures ANOVA (mixed effects model) with Tukey’s multiple comparisons. Data were considered statistically significant at *p*<0.05.

## Data availability

All data will be made publicly available upon publication and upon request for peer review.

## Acknowledgements

We thank Alice Mueller, Henning Jacobsen, Abhinaya Ganesan, Stephanie Peralta, and Sharvari Deshpande for their assistance with breeding, animal work, and some assays. We thank Art Arnold at UCLA for providing transgenic male FCG mice for breeding at Johns Hopkins. We are grateful to members of the Davis, Klein, and Pekosz labs at Johns Hopkins Bloomberg School of Public Health for their feedback on this work.

## Author contributions

Santosh Dhakal, Robert Maul, and Sabra Klein conceived the experimental questions contained in the funded NIH application. Santosh Dhakal, Han-Sol Park, Kumba Seddu, and Patrick Creisher conducted all mouse work. Santosh Dhakal, Kumba Seddu, and John Lee conducted antibody assays and Kumba Seddu and Patrick Creisher conducted steroid assays. Santosh Dhakal, Han-Sol Park, and Kumba Seddu conducted all flow cytometry and Santosh Dhakal and Han-Sol Park did all flow cytometry analyses. Robert Maul and Isabella Hernandez conducted FACS sorting of germinal center B cells. Han-Sol Park and Kimberly Davis conducted analyses of germinal center B cells. Santosh Dhakal and Han-Sol Park organized all data, conducted all statistical analyses, and created all figures. Santosh Dhakal and Sabra Klein wrote the manuscript and all authors approved of the final draft.

## References

1. Engler RJ, Nelson MR, Klote MM, VanRaden MJ, Huang CY, Cox NJ, Klimov A, Keitel WA, Nichol KL, Carr WW, Treanor JJ. 2008. Half-vs full-dose trivalent inactivated influenza vaccine (2004-2005): age, dose, and sex effects on immune responses. Arch Intern Med 168:2405–14.

2. Furman D, Hejblum BP, Simon N, Jojic V, Dekker CL, Thiébaut R, Tibshirani RJ, Davis MM. 2014. Systems analysis of sex differences reveals an immunosuppressive role for testosterone in the response to influenza vaccination. Proc Natl Acad Sci U S A 111:869–74.

3. Potluri T, Fink AL, Sylvia KE, Dhakal S, Vermillion MS, Vom Steeg L, Deshpande S, Narasimhan H, Klein SL. 2019. Age-associated changes in the impact of sex steroids on influenza vaccine responses in males and females. NPJ Vaccines 4:29.

4. Fink AL, Engle K, Ursin RL, Tang WY, Klein SL. 2018. Biological sex affects vaccine efficacy and protection against influenza in mice. Proc Natl Acad Sci U S A 115:12477–12482.

5. Živković I, Petrović R, Arsenović-Ranin N, Petrušić V, Minić R, Bufan B, Popović O, Leposavić G. 2018. Sex bias in mouse humoral immune response to influenza vaccine depends on the vaccine type. Biologicals 52:18–24.

6. Ursin RL, Dhakal S, Liu H, Jayaraman S, Park HS, Powell HR, Sherer ML, Littlefield KE, Fink AL, Ma Z, Mueller AL, Chen AP, Seddu K, Woldetsadik YA, Gearhart PJ, Larman HB, Maul RW, Pekosz A, Klein SL. 2022. Greater Breadth of Vaccine-Induced Immunity in Females than Males Is Mediated by Increased Antibody Diversity in Germinal Center B Cells. mBio 13:e0183922.

7. Frasca D, Blomberg BB, Garcia D, Keilich SR, Haynes L. 2020. Age-related factors that affect B cell responses to vaccination in mice and humans. Immunol Rev 296:142–154.

8. Saurwein-Teissl M, Lung TL, Marx F, Gschösser C, Asch E, Blasko I, Parson W, Böck G, Schönitzer D, Trannoy E, Grubeck-Loebenstein B. 2002. Lack of antibody production following immunization in old age: association with CD8(+)CD28(-) T cell clonal expansions and an imbalance in the production of Th1 and Th2 cytokines. J Immunol 168:5893–9.

9. Shapiro JR, Li H, Morgan R, Chen Y, Kuo H, Ning X, Shea P, Wu C, Merport K, Saldanha R, Liu S, Abrams E, Chen Y, Kelly DC, Sheridan-Malone E, Wang L, Zeger SL, Klein SL, Leng SX. 2021. Sex-specific effects of aging on humoral immune responses to repeated influenza vaccination in older adults. npj Vaccines 6:147.

10. Dhakal S, Deshpande S, McMahon M, Strohmeier S, Krammer F, Klein SL. 2022. Female-biased effects of aging on a chimeric hemagglutinin stalk-based universal influenza virus vaccine in mice. Vaccine 40:1624–1633.

11. Ferdinands JM, Gaglani M, Martin ET, Monto AS, Middleton D, Silveira F, Talbot HK, Zimmerman R, Patel M. 2021. Waning Vaccine Effectiveness Against Influenza-Associated Hospitalizations Among Adults, 2015–2016 to 2018–2019, United States Hospitalized Adult Influenza Vaccine Effectiveness Network. Clinical Infectious Diseases 73:726-729.

12. Hu W, Sjoberg PA, Fries AC, DeMarcus LS, Robbins AS. 2022. Waning Vaccine Protection against Influenza among Department of Defense Adult Beneficiaries in the United States, 2016-2017 through 2019-2020 Influenza Seasons. Vaccines (Basel) 10.

13. Hsu JP, Zhao X, Chen MIC, Cook AR, Lee V, Lim WY, Tan L, Barr IG, Jiang L, Tan CL, Phoon MC, Cui L, Lin R, Leo YS, Chow VT. 2014. Rate of decline of antibody titers to pandemic influenza A (H1N1-2009) by hemagglutination inhibition and virus microneutralization assays in a cohort of seroconverting adults in Singapore. BMC Infectious Diseases 14:414.

14. Pone EJ, Zhang J, Mai T, White CA, Li G, Sakakura JK, Patel PJ, Al-Qahtani A, Zan H, Xu Z, Casali P. 2012. BCR-signalling synergizes with TLR-signalling for induction of AID and immunoglobulin class-switching through the non-canonical NF-κB pathway. Nature Communications 3:767.

15. Castiblanco DP, Maul RW, Russell Knode LM, Gearhart PJ. 2017. Co-Stimulation of BCR and Toll-Like Receptor 7 Increases Somatic Hypermutation, Memory B Cell Formation, and Secondary Antibody Response to Protein Antigen. Front Immunol 8:1833.

16. Cunningham MA, Wirth JR, Naga O, Eudaly J, Gilkeson GS. 2014. Estrogen Receptor Alpha Binding to ERE is Required for Full Tlr7-and Tlr9-Induced Inflammation. SOJ Immunol 2.

17. Pauklin S, Sernández IV, Bachmann G, Ramiro AR, Petersen-Mahrt SK. 2009. Estrogen directly activates AID transcription and function. J Exp Med 206:99–111.

18. Dodd KC, Menon M. 2022. Sex bias in lymphocytes: Implications for autoimmune diseases. Front Immunol 13:945762.

19. Pollard AJ, Bijker EM. 2021. A guide to vaccinology: from basic principles to new developments. Nat Rev Immunol 21:83–100.

20. Harrell MI, Iritani BM, Ruddell A. 2008. Lymph node mapping in the mouse. J Immunol Methods 332:170–4.

21. Burgoyne PS, Arnold AP. 2016. A primer on the use of mouse models for identifying direct sex chromosome effects that cause sex differences in non-gonadal tissues. Biol Sex Differ 7:68.

22. Krammer F. 2019. The human antibody response to influenza A virus infection and vaccination. Nature Reviews Immunology 19:383–397.

23. Song JY, Cheong HJ, Hwang IS, Choi WS, Jo YM, Park DW, Cho GJ, Hwang TG, Kim WJ. 2010. Long-term immunogenicity of influenza vaccine among the elderly: Risk factors for poor immune response and persistence. Vaccine 28:3929–3935.

24. Frasca D, Blomberg BB. 2020. Aging induces B cell defects and decreased antibody responses to influenza infection and vaccination. Immunity & Ageing 17:37.

25. Mesin L, Ersching J, Victora GD. 2016. Germinal Center B Cell Dynamics. Immunity 45:471–482.

26. Maul RW, Gearhart PJ. 2010. AID and somatic hypermutation. Adv Immunol 105:159–91.

27. Sasaki S, Sullivan M, Narvaez CF, Holmes TH, Furman D, Zheng NY, Nishtala M, Wrammert J, Smith K, James JA, Dekker CL, Davis MM, Wilson PC, Greenberg HB, He XS. 2011. Limited efficacy of inactivated influenza vaccine in elderly individuals is associated with decreased production of vaccine-specific antibodies. J Clin Invest 121:3109–19.

28. Henry C, Zheng NY, Huang M, Cabanov A, Rojas KT, Kaur K, Andrews SF, Palm AE, Chen YQ, Li Y, Hoskova K, Utset HA, Vieira MC, Wrammert J, Ahmed R, Holden-Wiltse J, Topham DJ, Treanor JJ, Ertl HC, Schmader KE, Cobey S, Krammer F, Hensley SE, Greenberg H, He XS, Wilson PC. 2019. Influenza Virus Vaccination Elicits Poorly Adapted B Cell Responses in Elderly Individuals. Cell Host Microbe 25:357–366.e6.

29. Fish EN. 2008. The X-files in immunity: sex-based differences predispose immune responses. Nat Rev Immunol 8:737–44.

30. Souyris M, Cenac C, Azar P, Daviaud D, Canivet A, Grunenwald S, Pienkowski C, Chaumeil J, Mejía J, Guéry J-C. 2018. TLR7 escapes X chromosome inactivation in immune cells. Science Immunology 3:eaap8855.

31. Klein SL, Flanagan KL. 2016. Sex differences in immune responses. Nat Rev Immunol 16:626–38.

32. vom Steeg LG, Vermillion MS, Hall OJ, Alam O, McFarland R, Chen H, Zirkin B, Klein SL. 2016. Age and testosterone mediate influenza pathogenesis in male mice. Am J Physiol Lung Cell Mol Physiol 311:L1234–L1244.

33. Robinson DP, Lorenzo ME, Jian W, Klein SL. 2011. Elevated 17β-estradiol protects females from influenza A virus pathogenesis by suppressing inflammatory responses. PLoS Pathog 7:e1002149.

34. Nguyen DC, Masseoud F, Lu X, Scinicariello F, Sambhara S, Attanasio R. 2011. 17β-Estradiol restores antibody responses to an influenza vaccine in a postmenopausal mouse model. Vaccine 29:2515–8.

35. Hill L, Jeganathan V, Chinnasamy P, Grimaldi C, Diamond B. 2011. Differential roles of estrogen receptors α and β in control of B-cell maturation and selection. Mol Med 17:211–20.

36. Park SR, Zan H, Pal Z, Zhang J, Al-Qahtani A, Pone EJ, Xu Z, Mai T, Casali P. 2009. HoxC4 binds to the promoter of the cytidine deaminase AID gene to induce AID expression, class-switch DNA recombination and somatic hypermutation. Nat Immunol 10:540–50.

37. Mai T, Zan H, Zhang J, Hawkins JS, Xu Z, Casali P. 2010. Estrogen receptors bind to and activate the HOXC4/HoxC4 promoter to potentiate HoxC4-mediated activation-induced cytosine deaminase induction, immunoglobulin class switch DNA recombination, and somatic hypermutation. J Biol Chem 285:37797–810.

38. Wilhelmson AS, Lantero Rodriguez M, Stubelius A, Fogelstrand P, Johansson I, Buechler MB, Lianoglou S, Kapoor VN, Johansson ME, Fagman JB, Duhlin A, Tripathi P, Camponeschi A, Porse BT, Rolink AG, Nissbrandt H, Turley SJ, Carlsten H, Mårtensson I-L, Karlsson MCI, Tivesten Å. 2018. Testosterone is an endogenous regulator of BAFF and splenic B cell number. Nature Communications 9:2067.

39. Vom Steeg LG, Flores-Garcia Y, Zavala F, Klein SL. 2019. Irradiated sporozoite vaccination induces sex-specific immune responses and protection against malaria in mice. Vaccine 37:4468–4476.

40. Kasmani MY, Topchyan P, Brown AK, Brown RJ, Wu X, Chen Y, Khatun A, Alson D, Wu Y, Burns R, Lin CW, Kudek MR, Sun J, Cui W. 2023. A spatial sequencing atlas of age-induced changes in the lung during influenza infection. Nat Commun 14:6597.

41. Henry C, Palm AE, Krammer F, Wilson PC. 2018. From Original Antigenic Sin to the Universal Influenza Virus Vaccine. Trends Immunol 39:70–79.

42. Dhakal S, Klein SL. 2019. Host Factors Impact Vaccine Efficacy: Implications for Seasonal and Universal Influenza Vaccine Programs. J Virol 93.

43. Arnold AP, Chen X. 2009. What does the “four core genotypes” mouse model tell us about sex differences in the brain and other tissues? Front Neuroendocrinol 30:1–9.

44. De Vries GJ, Rissman EF, Simerly RB, Yang LY, Scordalakes EM, Auger CJ, Swain A, Lovell-Badge R, Burgoyne PS, Arnold AP. 2002. A model system for study of sex chromosome effects on sexually dimorphic neural and behavioral traits. J Neurosci 22:9005–14.

